# CrkL is required for donor T cell migration to GvHD Target organs

**DOI:** 10.1101/854042

**Authors:** Nathan H. Roy, Mahinbanu Mammadli, Janis K. Burkhardt, Mobin Karimi

**Author notes:** Author for correspondence: Mobin Karimi, Assistant Professor of Immunology and Microbiology, SUNY Upstate Medical University, 766 Irving Ave Weiskotten Hall Suite 2281, Syracuse, NY 13210.

## Abstract

The success of cancer therapies based on allogeneic hematopoietic stem cell transplant relies on the ability to separate graft-versus-host disease (GvHD) from graft-versus-tumor (GVT) responses. Controlling donor T cell migration into peripheral tissues is a viable option to limit unwanted tissue damage, but a lack of specific targets limits progress on this front. Here, we show that the adaptor protein CrkL, but not the closely related family members CrkI or CrkII, is a crucial regulator of T cell migration. *In vitro*, CrkL-deficient T cells fail to polymerize actin in response to the integrin ligand ICAM-1, resulting in defective migration. Using a mouse model of GvHD/GVT, we found that while CrkL-deficient T cells can efficiently eliminate hematopoietic tumors they are unable to migrate into inflamed organs, such as the liver and small intestine, and thus do not cause GvHD. These results suggest a specific role for CrkL in trafficking to peripheral organs but not the lymphatic system. In line with this, we found that although CrkL-deficient T cells could clear hematopoietic tumors, they failed to clear the same tumor growing subcutaneously, highlighting the role of CrkL in controlling T cell migration into peripheral tissues. Our results define a unique role for CrkL in controlling T cell migration, and suggest that CrkL function could be therapeutically targeted to enhance the efficacy of immunotherapies involving allogeneic donor cells.

## INTRODUCTION

T cell migration out of the vasculature into peripheral tissue is a key control point in the inflammatory response. Although trafficking to tissues is required for protective immunity, uncontrolled T cell infiltration into tissue can result in exacerbated inflammation and tissue damage. T cell recruitment is highly regulated by the local endothelial cells which, in response to pro-inflammatory cues, up-regulate molecules such as chemokines and adhesion receptors. These molecules promote T cell adhesion to the endothelial monolayer, and guide migration through the monolayer into the tissue ^1^. The endothelial ligands ICAM-1 and VCAM-1 are recognized by the T cell integrins LFA-1 and VLA-4, respectively, which mediate firm adhesion to the vascular wall and subsequent migration into inflamed tissue ^1, 2^. In addition to acting as adhesion receptors, integrins also initiate downstream signaling events that are important for T cell migration. For instance, ligation of LFA-1 immediately results in T cell polarization, actin polymerization, and migration, all of which are necessary for crossing the endothelial barrier ^3, 4, 5, 6, 7^. Integrin blocking antibodies have been used successfully to treat multiple sclerosis and other inflammatory diseases, but this approach interferes with a broad range of responses. Disruption of signaling events downstream of engaged integrins could yield more context-specific therapies, but to develop such strategies, a better understanding of the relevant signaling pathways is required.

We recently identified the CT10 Regulator of Kinase (Crk) family of adapter proteins as crucial intermediaries that link LFA-1 engagement to actin responses and migration in T cells ^7^. Activated T cells lacking Crk proteins fail to polymerize actin downstream of LFA-1, and migrate slower and less directionally than their WT counterparts ^7^. Importantly, these mutant T cells also have defects in crossing endothelial monolayers *in vitro*, and they fail to traffic to sites of inflammation *in vivo* ^8^. Additionally, using a clinically relevant mouse model of graft-versus-host disease / graft-versus-leukemia (GvHD/GVL), we showed that T cells lacking Crk proteins can efficiently clear tumor cells but cause little-to-no GvHD pathology ^8^. Consistent with the view that Crk proteins mediate integrin-dependent trafficking in a tissue-specific manner, we found that Crk deficient T cells could not migrate to the target GvHD organs liver and small intestine (SI), although they trafficked efficiently to the secondary lymphoid organs spleen and lymph nodes. These findings identify Crk proteins as potential targets for tissue-selective disruption of integrin-dependent inflammatory responses.

Development of Crk proteins as therapeutic targets requires addressing redundancy among family members. The Crk family consists of three isoforms transcribed from two loci. CrkI and CrkII are transcribed from the *crk* locus, while the paralog CrkL is transcribed from the *crkl* locus. These proteins are composed of a single N-terminal SH2 domain, followed by either two consecutive SH3 domains (CrkII and CrkL), or a single SH3 domain (CrkI) ^9^. The Crk proteins are widely expressed across tissues and have many biological functions, all of which stem from their role as adaptor proteins that coordinate signaling complexes downstream of cell surface receptors ^9, 10^. Crk proteins are particularly important for adhesion and migration ^11^. They have been shown to localize to adhesion sites and regulate the stability of these structures in non-hematopoietic cells ^12, 13, 14, 15^, and alterations in their expression is associated with invasive potential in several tumors ^16, 17, 18^. Crk family members share many of the same binding partners, and they have been shown to have overlapping functions in some processes ^15, 19, 20^. On the other hand, there are clear instances (most notably during development) where the Crk proteins have non-overlapping roles, showing that in some instances they have evolved separate functions ^21, 22^. All of our previous work defining the role of Crk proteins in T cell migration was performed using a mouse floxed for both the *crk* and *crkl* loci and crossed with a CD4-cre mouse, resulting in T cells devoid for all three family members (herein referred to as DKO). Therefore, it is unclear if the Crk family members work together to promote T cell migration, or if only a single Crk isoform is responsible for this function. Now, using T cells lacking either CrkI/II or CrkL, we show that CrkL is the dominant Crk family member that controls T cell migration. T cells lacking CrkI/II show a WT phenotype, whereas T cells lacking CrkL phenocopy DKO T cells in their responses to ICAM-1 *in vitro* and *in vivo* in a GvHD/GVT mouse model. This work has defined a unique role for CrkL in T cell migration, opening the door to novel therapeutic approaches based on targeting CrkL function.

## RESULTS

### CrkL is needed for T cell spreading and migration in response to the integrin ligand ICAM-1

We showed previously that T cells lacking all Crk family members exhibit defects in integrin-dependent migration and *in vivo* trafficking ^7, 8^. To ask if this aspect of Crk protein function depends on a single protein isoform or if there is functional redundancy among family members, we used mice that are floxed for either the *crk* or the *crkL* locus, crossed with CD4-Cre mice to specific delete these loci in T cells. The resulting T cells are lacking either CrkI/II or CrkL (**Fig 1A**). We first tested the ability of these T cells to polymerize actin and migrate in response to surface-presented integrin ligands *in vitro*. Rested CD4^+^ T cell blasts were prepared as described in Materials and Methods, and then allowed to interact with ICAM-1 coated surfaces, fixed, and labelled with phalloidin to visualize F-actin. As shown previously ^4, 5, 7^, WT T cells adhered tightly to the surfaces, and formed a polarized morphology characteristic of cells migrating on ICAM-1 (**Fig 1B**). These WT cells showed strong actin polymerization, especially at the leading edge (**Fig 1B**). T cells lacking CrkI/II were indistinguishable from the WT cells. In contrast, DKO and CrkL KO T cells failed to spread, and lacked a clear F-actin-rich leading edge (**Fig 1B**). Quantitation of the total F-actin signal confirmed that DKO and CrkL KO T cells responding to ICAM-1 contained significantly less F-actin than WT or CrkI/II KO T cells (**Fig 1C**). To determine if the defects in actin responses result in altered migration, we imaged T cells migrating on ICAM-1 coated surfaces and tracked cell movement. DKO and CrkL KO T cells migrated significantly slower than WT or CrkI/II KO cells (**Fig 1D**). As we showed previously for DKO T cells ^7^, loss of a clear actin-rich leading edge also resulted in reduced directional persistence in CrkL KO T cells (data not shown). Together, these data show that the two products of the Crk locus, CrkI and CrkII, are dispensable for actin polymerization and migration downstream of integrin engagement. The CrkL protein is the relevant family member controlling these events in T cells.

**Figure 1.**
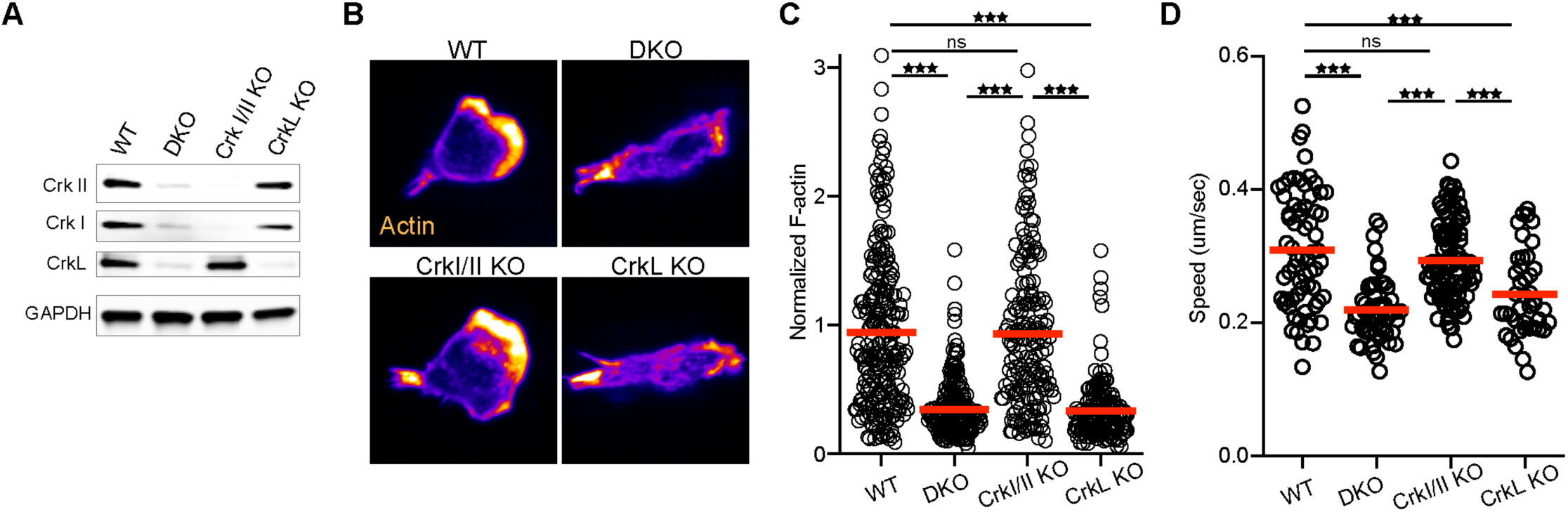
CrkL is required for T cell integrin responses. **A)** Western blot of T cells purified from WT or knockout mice. **B)** Activated T cells from the indicated genotypes were allowed to settle on ICAM-1 coated coverslips, fixed, and stained for F-actin using fluorescent phalloidin. **C)** Quantification of total F-actin signal from **(B)**, pooled from n=3. **D)** Activated T cells from the indicated genotypes were imaged migrating on ICAM-1 coated surfaces and average speed was calculated, pooled from n=3. A one-way ANOVA was used to calculate p-values. *p<0.05; **p<0.01; ***p<0.001

### T cells lacking CrkL clear hematopoietic tumors but fail to traffic to target GvHD organs

To ask if CrkL is also the critical Crk family member responsible for T cell migration *in vivo*, we utilized a graft-versus-host / graft-versus-tumor mouse model (GvHD/GVT). In this model, GvHD occurs in a subset of organs and involves early migration of alloreactive T cells into these organs followed by T cell expansion and later tissue destruction ^23^. To test the role of the individual Crk proteins in GvHD pathology, lethally irradiated BALB/c mice were injected with T-cell depleted C57BL/6 bone marrow (TCDBM) along with CD8^+^ T cells from WT, DKO, CrkI/II KO, or CrkL KO C57BL/6 mice. In addition, A20 lymphoma cells expressing luciferase were also injected, to determine if the transferred CD8^+^ T cells could eliminate tumor cells. Mouse weights, GvHD clinical score, tumor burden, and overall survival were monitored. As expected, transfer of WT CD8^+^ T cells caused severe GvHD with no detectable signs of tumors at the time of sacrifice (**Fig 2A-D**). As we showed previously ^8^, DKO CD8^+^ T cells did not cause GvHD, but still efficiently cleared the tumor cells, resulting in healthy mice throughout the course of the experiment. CrkL KO CD8^+^ T cells recapitulated the DKO phenotype, while CrkI/II KO CD8^+^ T cells had a phenotype indistinguishable from the WT. This pattern was observed for all disease metrics, including overall survival (**Fig 2A-D**).

**Figure 2.**
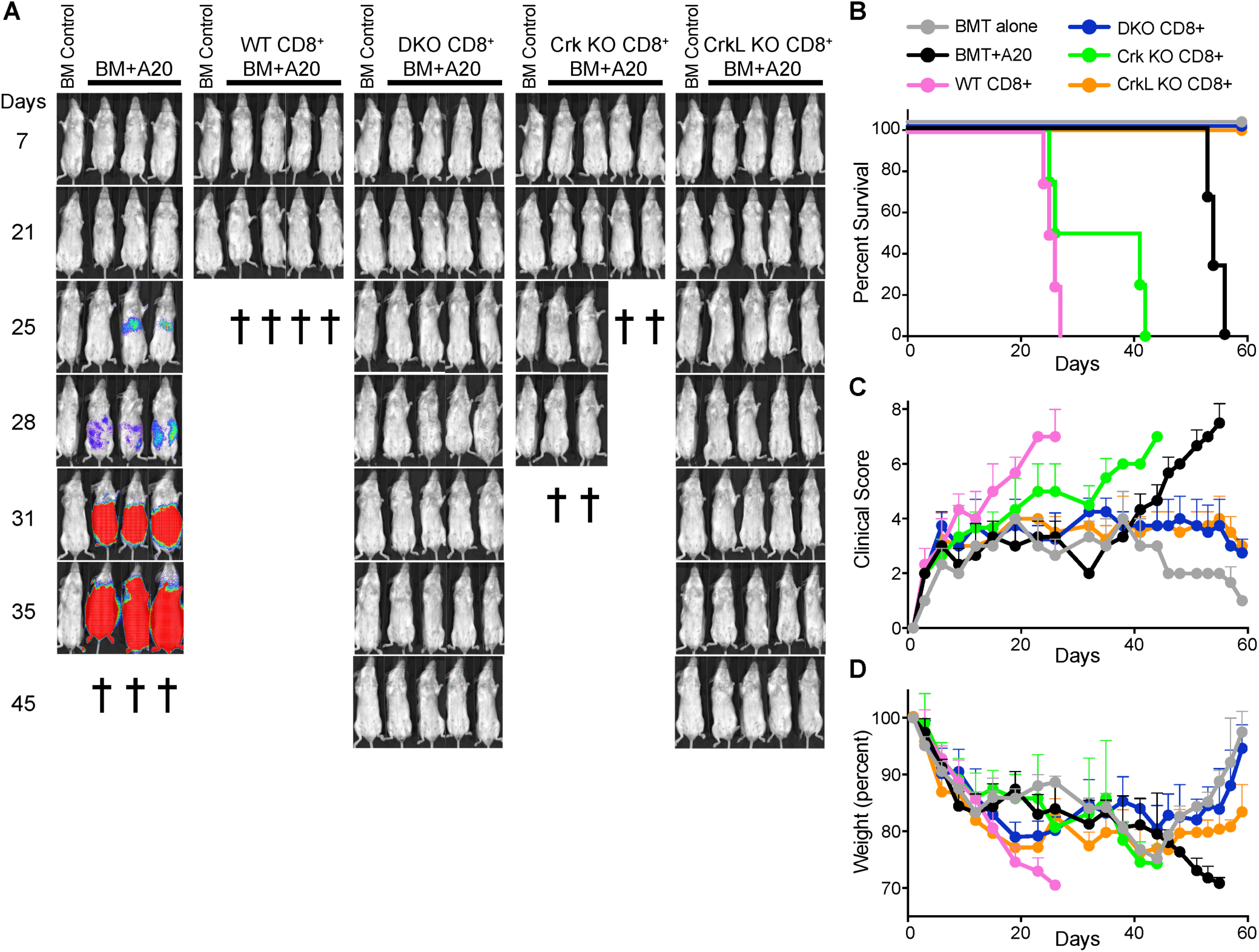
T cells lacking CrkL do not cause pathology in a model of GvHD/GVL. Lethally irradiated BALB/c mice were injected with T-cell depleted C57BL/6 bone marrow along with A20 lymphoma cells expressing luciferase. At the same time, CD8^+^ T cells from WT, DKO, CrkI/II KO, or CrkL KO mice were also injected. Over the next several weeks, the mice were periodically injected with luciferin and imaged to visualize tumor burden. **A)** Images of individual mice, **B)** Overall survival, **C)** clinical score, and **D)** mouse weights were recorded. Representative experiment of n=3

Based on the failure of CrkL KO T cells to migrate normally *in vitro*, we reasoned that CrkL KO T cells do not cause GvHD because of their inability to traffic to target GvHD organs. However, because the GvHD/GVT experiments are take place over the course of several weeks, it is difficult to establish a causative role for T cell migration in initiating the inflammatory response. We therefore set up a short-term competitive experiment to examine the role of Crk proteins in T cell migration into target GvHD organs. Lethally irradiated BALB/c mice were injected with TCDBM along with a 1:1 mix of WT CD45.1^+^ CD8^+^ T cells (purified from B6.SJL mice) and either WT, DKO, CrkI/II KO, or CrkL KO CD8^+^ T cells, all of which are CD45.2^+^. Seven days post injection, before the onset of GvHD pathology, the mice were sacrificed and their livers, small intestines, and spleens were processed for flow cytometric analysis. As expected, when WT CD45.1^+^ T cells were mixed with control WT CD45.2^+^ T cells, the two were detected in approximately a 1:1 ratio in every organ tested (**Fig 3A-B**). This was also true for CrkI/II KO T cells, indicating that Crk I and Crk II are dispensable for T cell migration *in vivo*. However, when DKO or CrkL KO T cells were used, these T cells had a significant defect in migration to the liver and the small intestine compared to their CD45.1^+^ WT counterparts. Importantly, migration into the spleen was largely normal, indicating that DKO and CrkL KO T cells can still traffic efficiently to lymphoid organs (**Fig 3A-B**). Taken together with the data in Figures 1 and 2, these results demonstrate that CrkL is the relevant Crk family protein controlling T cell migration both *in vitro* and *in vivo*, and highlights the importance of CrkL-dependent T cell trafficking during inflammation and GvHD.

**Figure 3.**
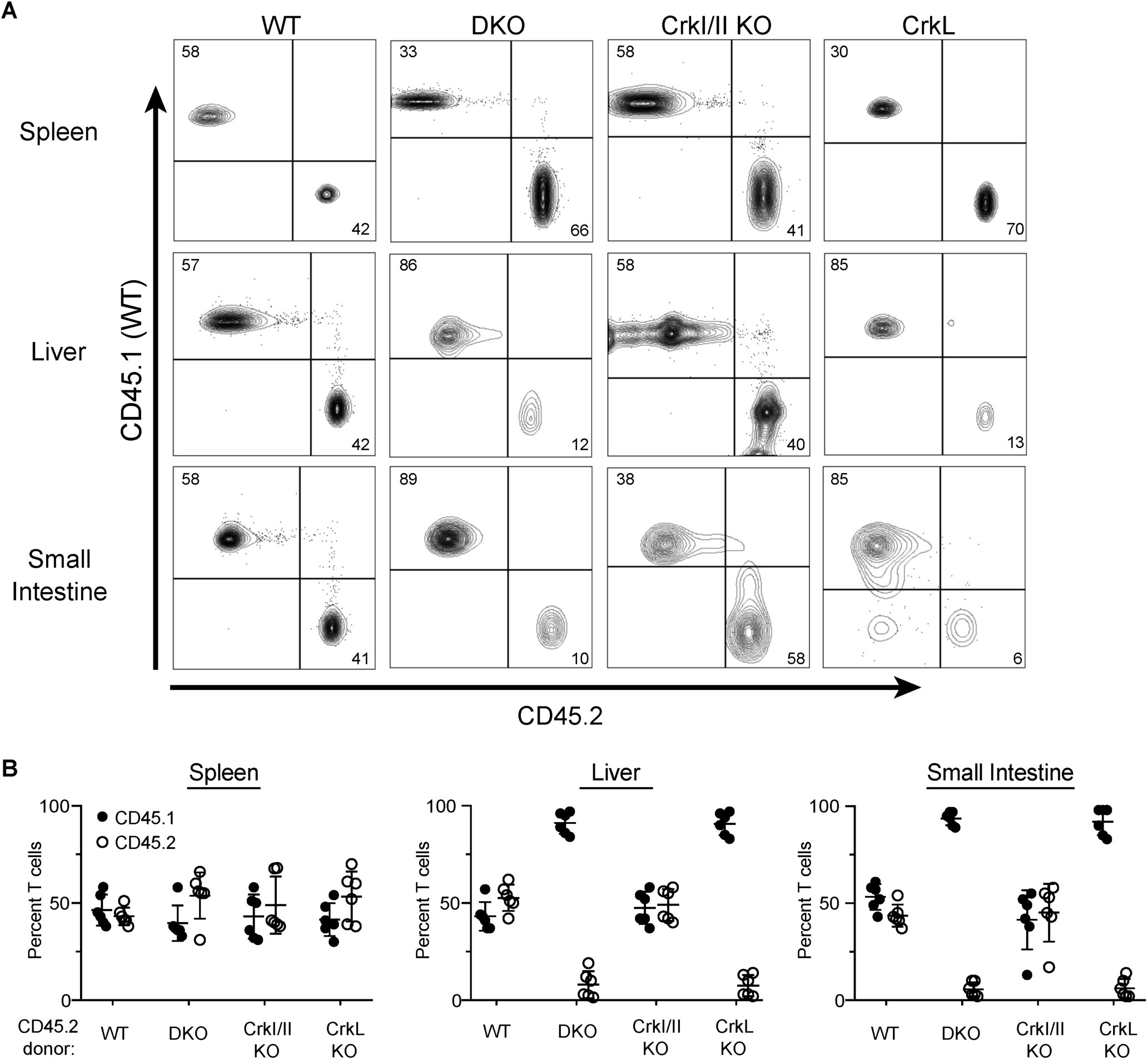
T cell trafficking to GvHD organs required CrkL. **A)** Lethally irradiated BALB/c mice were injected with T-cell depleted C57BL/6 bone marrow along with a 1:1 ratio of WT CD45.1^+^ CD8^+^ T cells with CD45.2^+^ CD8^+^ T cells from either WT, DKO, CrkI/II KO, or CrkL KO mice. Seven days post injection, the mice were sacrificed and the spleen, liver, and small intestines were processed for flow cytometry. Gating was done on donor CD8^+^ T cells, and the percentages of CD45.1^+^ and CD45.2^+^ T cells were determined. A single mouse from a representative experiment is shown. **B)** Quantification of CD45.1^+^ to CD45.2^+^ percentages from indicated tissues. n=3

### CrkL KO T cells fail to eliminate a sub-cutaneous tumor

The trafficking studies above suggest that the ability of CrkL KO T cells to clear lymphoma cells while inducing minimal GvHD pathology stems from their ability to migrate efficiently to lymphoid tissues where the tumor resides, but not to GvHD target organs. If so, one would predict that if the tumor were located in a peripheral tissue outside of the lymphatic system, CrkL KO T cells would not be able to access and kill the tumor. To test this, we set up a GvHD/GVT experiment similar to that in Figure 2, except that the A20 tumor cells were injected subcutaneously as opposed to intravenously. As shown in **Figure 4**, both WT and CrkI/II KO T cells efficiently cleared the subcutaneous tumor, but these animals succumbed to GvHD. In contrast, DKO and CrkL KO T cells were unable to clear the subcutaneous tumor and these recipients were sacrificed due to the resulting tumor pathology (**Fig 4A-D**). Together, these data indicate that deletion of CrkL in T cells can separate GvHD from GVT effects, but only for tumors that reside in the circulation and in secondary lymphoid organs (e.g. hematologic malignancies). More broadly, these data provide strong evidence that the CrkL controls T cell entry into inflamed tissues, and that disruption of CrkL function can be used to perturb this trafficking, while preserving trafficking to lymphoid organs.

**Figure 4.**
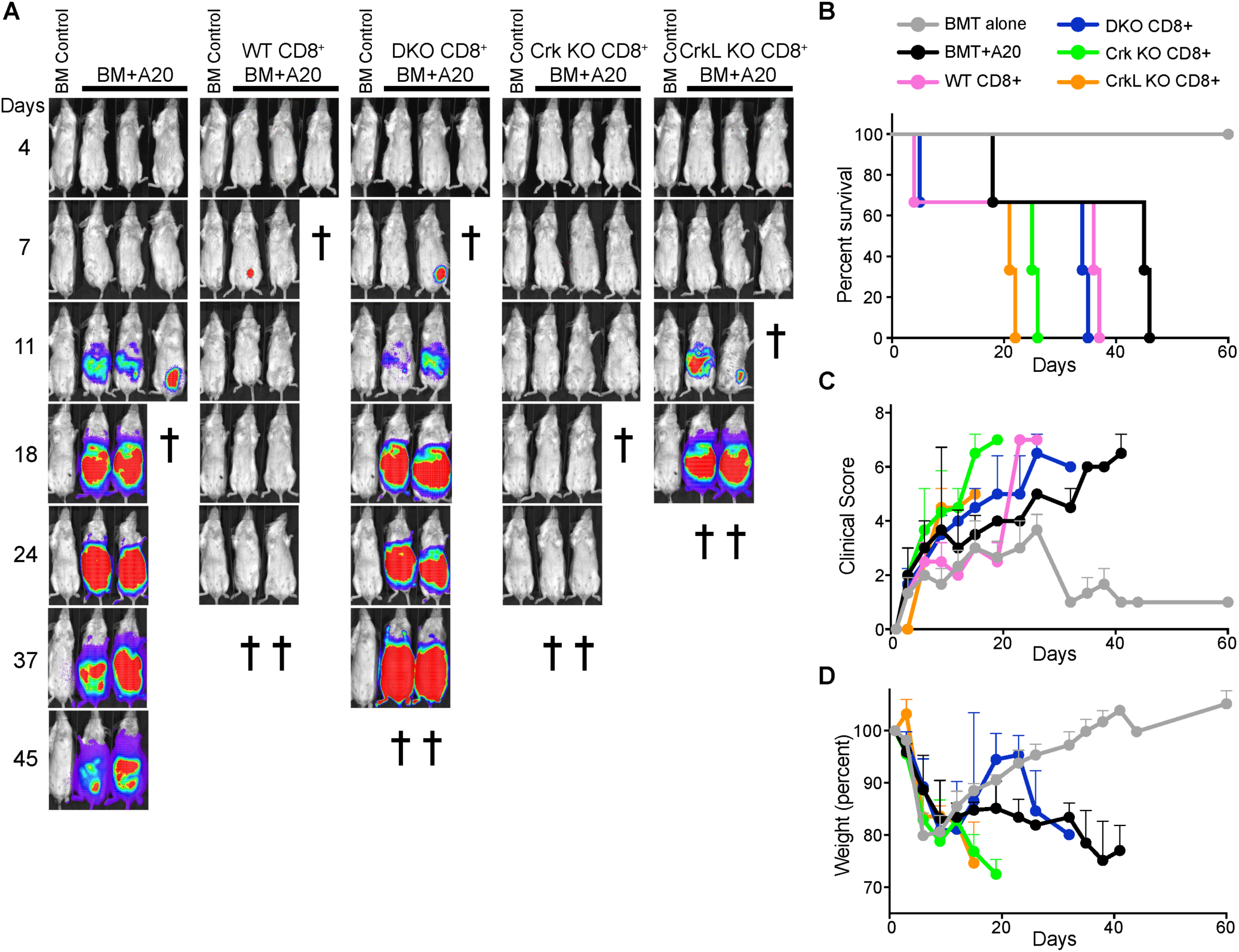
T cells lacking CrkL fail to eliminate a subcutaneous tumor. **A)** Lethally irradiated BALB/c mice were injected with T-cell depleted C57BL/6 bone marrow along with CD8^+^ T cells from WT, DKO, CrkI/II KO, or CrkL KO mice. At the same time, A20 lymphoma cells expressing luciferase were injected subcutaneously. Over the next several weeks, the mice were periodically injected with luciferin and imaged to visualize tumor burden. **B)** Overall survival, **C)** clinical score, and **D)** mouse weights were recorded. Representative experiment of n=3

## DISCUSSION

We showed previously that T cells lacking all Crk family proteins exhibit defective migration into inflamed tissue *in vivo*, and an inability to carry out transendothelial migration (TEM) *in vitro* ^8^. Interestingly, this *in vitro* phenotype could be rescued by overexpression of any Crk family member ^8^, raising the possibility that Crk proteins might exhibit functional redundancy with respect to T cell migration. Here, we have carried out more stringent *in vitro* and *in vivo* analysis using single and double knockout T cells. We show that CrkL, but not CrkI or CrkII, is required for T cell actin polymerization and migration in response to ICAM-1. In an *in vivo* GvHD/GVT model, CrkL deficient T cells failed to traffic to GvHD target organs and did not cause inflammation, but could efficiently clear hematopoietic tumor cells. However, CrkL KO T cells could not clear the same tumor when it was growing subcutaneously, highlighting the role of CrkL in controlling T cell migration into peripheral tissues. Taken together, our studies identify a unique role for CrkL in T cell migration *in vitro* and *in vivo*, opening the door for future studies aimed at disrupting CrkL biology to alter T cell function.

Crk proteins have long been associated with adhesion and migration in many cell types including fibroblasts, hematopoietic cell lines, and cancer cells ^10, 11, 18^. They fulfill this function by linking tyrosine phosphorylated proteins to effector proteins that govern adhesion and migration, such as Src and Abl family kinases, GEFs and GAPs ^9, 10^. In this way, Crk proteins coordinate multi-protein complexes downstream of cell surface receptors. In T cells, only a small number of proteins are known to interact with Crk proteins ^24^. Based on our data defining a clear role for CrkL in actin polymerization and T cell migration, it is likely that CrkL has unique binding partners in T cells that mediate these phenotypes. The isolated SH2 and SH3 domains of CrkII and CrkL are very similar and have nearly identical binding affinities for protein ligands ^25^. Nonetheless, the full-length proteins show significant differences. In the intact protein, the SH2 domain of CrkII binds efficiently to tyrosine phosphorylated ligands, whereas the SH2 domain of CrkL shows low binding activity. Conversely, the N-terminal SH3 domain of CrkL binds freely to proline-rich peptides, but this activity is largely masked in CrkII ^25^. Further differences are seen upon tyrosine phosphorylation of Crk proteins, where the phosphorylation of CrkII (but not CrkL) further reduces binding to proline-rich ligands. Ligand binding of CrkII is also regulated by proline isomerization ^26, 27, 28^. Since the relevant proline is absent from CrkL, this likely represents another structural difference that regulates isoform-specific binding. Future work aimed at determining the specific binding partners of the different Crk proteins in primary T cells under different phosphorylation and isomerization conditions will be crucial for formulating therapies that alter CrkL dependent signaling pathways to tune T cell function.

We show that CrkL KO T cells fail to enter inflamed tissues and subcutaneous tumors, but they efficiently access lymphoid tissues. The selective requirement for CrkL in entering inflamed tissue likely reflects the requisite role of CrkL in regulating actin dynamics downstream of integrin engagement. T cells actively migrate on inflamed postcapillary venules and make numerous actin-dependent invadopodia-like protrusions to push their way through the endothelial monolayer ^6, 29, 30^. Therefore, the ability of the T cell to polymerize actin, migrate, and probe the endothelium may be defining factors for the ability to cross postcapillary venules. The preferential ability of CrkL KO T cells to access lymph nodes is likely to be attributable to differences in endothelial barriers. To access lymph nodes, T cells must cross high endothelial venules, specialized structures that create large pockets to fully engulf transmigrating T cells, efficiently escorting them across the barrier ^31^. This process may be much less demanding with respect to force generation by the T cell actin cytoskeleton.

Our studies showing that CrkL mediates integrin responses in migrating T cells expands upon earlier work showing that Crk proteins are important T cell activation. In that setting, CrkI/II are the best studied family members, but functional redundancy has not been systematically addressed. In particular, CrkII has been documented to bind a handful of proteins downstream of the T cell receptor, including ZAP-70, cCbl, CasL, and even the CD3ζ chain ^32, 33, 34, 35, 36^. Interestingly, both CrkII and CrkL have been implicated in LFA-1 activation at the immunological synapse, but they seem to achieve this via different mechanisms. Dephosphorylation of CrkII by SHP-1 controls CrkII’s ability to activate Rap1 in the integrin-rich zone of the synapse ^37^. On the other hand, CrkL regulates integrin activation by pathway involving the actin nucleator WAVE2 ^38^. In addition, CrkL has also been implicated in regulating WASp activation downstream of T cell receptor engagement ^39^. These two studies showing that CrkL regulates actin function at the IS, together with the work descried here showing a role for CrkL in actin regulation during T cell migration, raise the interesting possibility that CrkL has specialized for regulating cytoskeletal function. Indeed, there have been no reports of CrkI or CrkII regulating actin dynamics in T cells, lending support to the idea that CrkL has evolved for regulating actin dynamics in T cells.

Going forward, it will be interesting to identify CrkL binding partners and other signaling molecules needed for integrin-dependent actin responses and passage into inflamed tissues. Targeting these molecules could be used to more finely control T cell trafficking in the treatment of inflammatory diseases and for the design of next-generation adoptive T cell therapies

## MATERIALS AND METHODS

### Antibodies and Reagents

Anti-CD3 (2C11) and anti-CD28 (PV1) were from BioXCell. Anti-CrkL (SC-319) was from Santa Cruz. Anti-CD3-FITC was from E-biosciences. Anti-Crk (#610036), Anti-CD8-FITC, anti-CD3, anti-CD45.1-Pacific Blue, anti-H-2Kb-Pacific Blue, anti-CD45.2-APC and LIVE/DEAD Fixable Aqua Dead Cell Stain Kit were from BD Bioscience. Secondary antibodies conjugated to appropriate fluorophores and AlexaFluor 488-conjugated phalloidin were obtained from Molecular Probes. Luciferin was purchased from Perkin Elmer and Gold Bio. Recombinant mouse ICAM-1-Fc was from R&D Systems.

### Mice and Cell Culture

Throughout this study, Crk fl/fl:CrkL fl/fl mice ^19^(C57BL/6 background) were used as WT. Mice lacking all Crk proteins in T cells (CD4 Cre:Crk fl/fl:CrkL fl/fl mice, DKO) were described previously ^7^. To generate single KO mice, Crk fl/fl:CrkL fl/fl mice were crossed with C57BL/6 mice. Resulting single Crk fl/fl or CrkL fl:fl mice were then crossed with CD4^+^ Cre mice, and offspring were used as a source of CrkI/II KO or CrkL KO T cells. BALB/c mice were used as recipients in the GvHD/GVL studies. Unless otherwise indicated, all tissue culture reagents were from Gibco. Primary mouse CD4^+^ T cells were purified from lymph nodes and spleens by negative selection using anti-MHCII and anti-CD8 hybridoma supernatants (M5/114 and 2.43, respectively) and anti-rat Ig magnetic beads (Qiagen BioMag). The resulting CD4^+^ T cells were then immediately activated on 24-well plates coated with anti-CD3 and anti-CD28 (1 ug/ml each) at 1×10^6^ cells per well. Activation was done in complete T cell media (DMEM, 5% FBS, penicillin/streptomycin, non-essential amino acids, Glutamax, and 55 μM β-mercaptoethanol). After 48h, cells were removed from activation and diluted 1:1 with complete T cell media containing recombinant IL-2 (25 U/mL final concentration, obtained through the NIH AIDS Reagent Program, Division of AIDS, NIAID, NIH from Dr. Maurice Gately, Hoffmann - La Roche Inc ^40^). T cells were used 5-6 days after activation.

### F-actin Quantification of Migrating T cells

Activated CD4^+^ T cells were resuspended in Leibovitz’s L-15 media (Gibco) supplemented with 2 mg/mL glucose and incubated at 37°C for 20 min. T cells were then added to ICAM-1 coated coverslips (coated with 2 *μ*g/mL overnight at 4°C). After 20 min at 37°C, cells were washed in L-15 and fixed in 3.7% paraformaldehyde in PBS. Cells were then blocked and permeabilized in PSG (PBS, 0.01% saponin, 0.05% fish skin gelatin) for 20 min, followed by 45 min with fluorescent phalloidin (Molecular Probes) in PSG. Cells were washed in PSG, mounted, and imaged using a 63x PlanApo 1.4 NA objective on an Axiovert 200M (Zeiss) with a spinning disk confocal system (Ultraview ERS6; PerkinElmer). Four z-planes spanning a total of 0.75µm were collected at the cell-surface interface with an Orca Flash 4.0 camera (Hamamatsu). Image analysis of the rendered stacks was conducted using Velocity v6.3 software. Cells were identified using the “Find Objects” command, using a low threshold on the actin channel, and total phalloidin staining was quantified per cell based on integrated pixel intensity.

### Quantification of Migration on ICAM-1

Lab-Tek 8 chamber slides (ThermoFisher) were coated with 2µg/mL ICAM-1 overnight at 4°C. Activated CD4^+^ T cells were washed and resuspended in L-15 containing 2mg/mL glucose. T cells were then added to the chambers, incubated 20 min, gently washed to remove all unbound cells, and imaged using a 10x phase contrast objective at 37°C on a Zeiss Axiovert 200M microscope equipped with an automated X-Y stage and a Roper EMCCD camera. Time lapse images were collected every 30 sec for 10 min using SlideBook 6 software (Intelligent Imaging Innovations). Movies were exported into ImageJ, and cells were tracked using the manual tracking plugin to calculate speed.

### Western Blotting

Proteins were separated using NuPAGE 4-12% BisTris gels, transferred to nitrocellulose membranes (0.45µm, BioRad) and blocked using LI-COR blocking buffer diluted 1:1 with PBS for 1 hr at RT. Primary antibodies were mixed in Tris-buffered saline, 0.1% tween-20 (TBST) with 2% BSA and membranes were probed overnight at 4°C. After washing, membranes were incubated with fluorescent secondary antibodies and imaged using a LI-COR Odyssey system.

### Graft-versus-Host Disease (GvHD) and Graft-versus-Tumor (GVT) studies

Lethally irradiated BALB/c mice (800 cGy) mice were injcted intravenously with 5×10^6^ T cell-depleted bone marrow (TCDBM) cells with or without 2×10^6^ FACS-sorted CD8^+^ T cells from either WT, DKO, CrkI/II KO or CrkL KO mice. For GVT experiments, A20 B lymphoma cells transduced with luciferase were cultured as described previously 41, and 1×10^5^ A20 cells were injected into the mice at the same time with TCDBM and CD8^+^ T cells. For skin tumor experiments, the A20 cells were injected subcutaneously 1×10^6^. Mice were evaluated twice a week from the time of tumor injection by bioluminescence imaging using the IVIS 200 Imaging System (Xenogen) as previously described 42. Clinical presentation of the mice was assessed 2-3 times per week using a scoring system that sums changes in 5 clinical parameters: weight loss, posture, activity, fur texture, and skin integrity 43. Mice were euthanized when their weight dropped to < 30% of their initial body weight.

For cell sorting to obtain purified CD8+ T cells, T cells were purified with anti-CD8 magnetic beads using MACS columns (Miltenyi Biotec, Auburn, CA) prior to cell surface staining. FACS sorting was performed with a FACSAria cell sorter (BD Biosciences). FACS-sorted populations were typically of >95% purity.

### In vivo migration assays

Lethally irradiated BALB/c mice were injected intravenously with 5×10^6^ WT T cell-depleted bone marrow (TCDBM). At the same time, FACS-sorted CD8^+^ T cells from CD45.1^+^ B6.SJL mice were mixed at a 1:1 ratio with CD8^+^ T cells from with WT, DKO, CrkI/II KO or CrkL KO CD45.2^+^ mice. Seven days post transplantation, the mice were sacrificed and lymphocytes from the liver, small intestine, spleen, and skin-draining lymph nodes were isolated. Livers were perfused with PBS, dissociated, and filtered with a 70 mm filter. The small intestine was washed in media, shaken in strip buffer at 37°C for 30 min to remove the epithelial cells, washed, digested with collagenase D (100 mg/ml), DNase (1mg/ml) for 30min in 37°C, and filtered with a 70 μm filter. Lymphocytes from the liver and intestines were further enriched using a 40% Percoll gradient. The cells were then analyzed for CD45.1^+^ and CD45.2^+^ CD8^+^ T cells by flow cytometry performed on an LSR-II or FACS Canto (BD Biosciences). Dead cells were excluded from analysis with LIVE/DEAD Fixable Aqua Dead Cell staining. Data were analyzed with FlowJo software (TreeStar, Ashland, OR).

### Statistical analysis

Statistics were calculated using GraphPad Prism 8. Student’s t-test was used to compare two groups; when more than two groups were compared, a one-way ANOVA was performed using multiple comparisons with a Tukey correction. *p<0.05; **p<0.01; ***p<0.001

## Acknowledgements

We thank members of the Burkhardt and Karimi laboratories for helpful discussions. This research was funded by grants from the National Blood Foundation Scholar Award to (MK) and the National Institutes of Health (NIH LRP #L6 MD0010106 and AI130182 to MK, AI120701 to JKB). NHR is supported by NIH T32 5T32AR007442-30, as well as previous support by a Cancer Research Institute Irvington Postdoctoral Fellowship.

## Author contributions

NHR and MM performed experiments; NHR, JKB, and MK designed experiments, analyzed the data, and wrote the manuscript.

